# High-throughput variant detection using a color-mixing strategy

**DOI:** 10.1101/2021.06.30.450651

**Authors:** Nina Guanyi Xie, Kerou Zhang, Ping Song, Renqiang Li, Junfeng Luo, David Yu Zhang

**Affiliations:** Department of Bioengineering, Rice University, Houston, TX, 77030, USA; Nuprobe China, 489 Zhengli Rd, Yangpu District, Shanghai, China

## Abstract

Many diseases are related to multiple genetic alterations along a single gene. Probing for highly multiple (>10) variants in a single qPCR tube is not possible due to a limited number of fluorescence channels and one variant per channel, so many more tubes are needed. Here, we experimentally validate our novel color-mixing strategy that uses fluorescence combinations as digital color codes to probe multiple variants simultaneously. The color-mixing strategy relies on a simple intra-tube assay that can probe for 15 variants as part of an inter-tube assay that can probe for an exponentially increased number of variants. The color-mixing strategy is achieved using multiplex double-stranded toehold probes modified with fluorophores and quenchers; the probes are designed to be quenched or luminous after binding to wildtype or variant templates. We used the color-mixing strategy to probe for 21 pathogenic mutations in thalassemia and to distinguish between heterozygous and homozygous variants in 6 tubes, with a specificity of 99% and a sensitivity of 94%. To support tuberculosis diagnosis, we used the same strategy to simultaneously probe in *Mycobacterium tuberculosis* for rifampicin-resistance mutations occurring within one 81-bp region and one 48-bp region in *rpoB* gene, plus five isoniazid-resistance mutations in *inhA* and *katG* genes.

## INTRODUCTION

Nucleic acid variants are important biomarkers for disease diagnosis. High-throughput detection of variants, including single nucleotide variants (SNVs), small insertions, and deletions (INDELs), can improve the sensitivity and specificity of diagnosis. Next-generation sequencing (NGS) is the most commonly used high-throughput variant detection method. However, the overall workflow of NGS is relatively long and the cost is fairly high [1]. Droplet digital polymerase chain reaction (ddPCR) is a highly sensitive detection method with a fast turnaround time that can be used to detect and quantitate variants accurately. However, due to the limitation of numbers of fluorescence channels used, ddPCR is usually used for single-plex or duplex variant detection.

Quantitative polymerase chain reaction (qPCR) is the most commonly used method because the instruments are easy to access, and the overall workflow is simple. qPCR can be used for detection and quantitation of single variants by TaqMan probes and Molecular Beacon [2]. However, the detection of highly multiplex variants is challenging in a qPCR reaction because the number of fluorescence channels available in a qPCR instrument is ≤ six [3]. With one channel used for each single-plex variant, simultaneous variant detection using a qPCR instrument is limited to 6-plex. Further, it is hard to distinguish a germline heterozygous variant (50%) from a homozygous variant (100%) using qPCR because the cycle threshold (Ct) value of a 50% variant sample can be close to that of a 100% variant sample.

Here we presented a color-mixing strategy that can be used to identify multiplex variants and their heterogeneity. The color-mixing strategy can be implemented through two steps. First, an asymmetric PCR amplification is performed to preferentially enrich one strand within the region of interest over the other; second, end-point fluorescence detection is enabled with multiplex rationally designed toehold probes [4, 5].

Using this strategy, we developed an assay that can detect the 21 most common pathogenic mutations in thalassemia and can further distinguish the heterogeneity of those mutations [6]. We also designed an assay for *Mycobacterium tuberculosis* bacteria (MTB) using this strategy that can theoretically detect any drug-resistant mutation within one 81-bp and one 48-bp region in the *rpoB* gene, which we experimentally validated for 10 mutations, and can also detect another 5 mutations in the *inhA* and *katG* genes [7-10].

## Materials and Methods Color-mixing strategy

### Intra-tube color-mixing design

Four commonly used fluorescence channels with minimum spectral overlap were used for each tube: ROX, CY5, FAM, and HEX. The final output from each fluorescence channel has 2 states, ‘On’ (1) and ‘Off’ (0). When making full use of the 4 fluorescence channels, there are 15 (2^4^ − 1) state combinations indicating 15 variants, excluding the all-off state. Each state combination was given a color code. Note that for this simple design, we expect that the genetic disorders this color-mixing strategy is targeting will be predominantly caused by single mutation in any individual; that is, there are no coexisting genetic variants within the genome, so the test sample would generally contain one variant. Additionally, this strategy would be suitable for infectious diseases in which the genetic mutations of pathogens cause drug resistance, assuming that there is usually single mutation within the gene that causes the drug resistance. In the case of multi-drug resistance, multiple genes would contribute to the multi-drug resistance, with each gene likely to have a single drug-resistance mutation.

Multiplex double-stranded toehold probes modified with 4 different fluorophores and quenchers were implemented in one tube. A single ‘On’ state from probes 1-4 can thus detect variants 1-4 (Figure 1A). Additionally, simultaneous signals from probes with different fluorophores can target other variants: two ‘On’ states from probe-5-ROX and probe-5-CY5 indicate the presence of variant 5; and three ‘On’ states from probe-14-CY5, probe-14-FAM and probe-14-HEX indicate the presence of variant 14.

**Figure 1.**
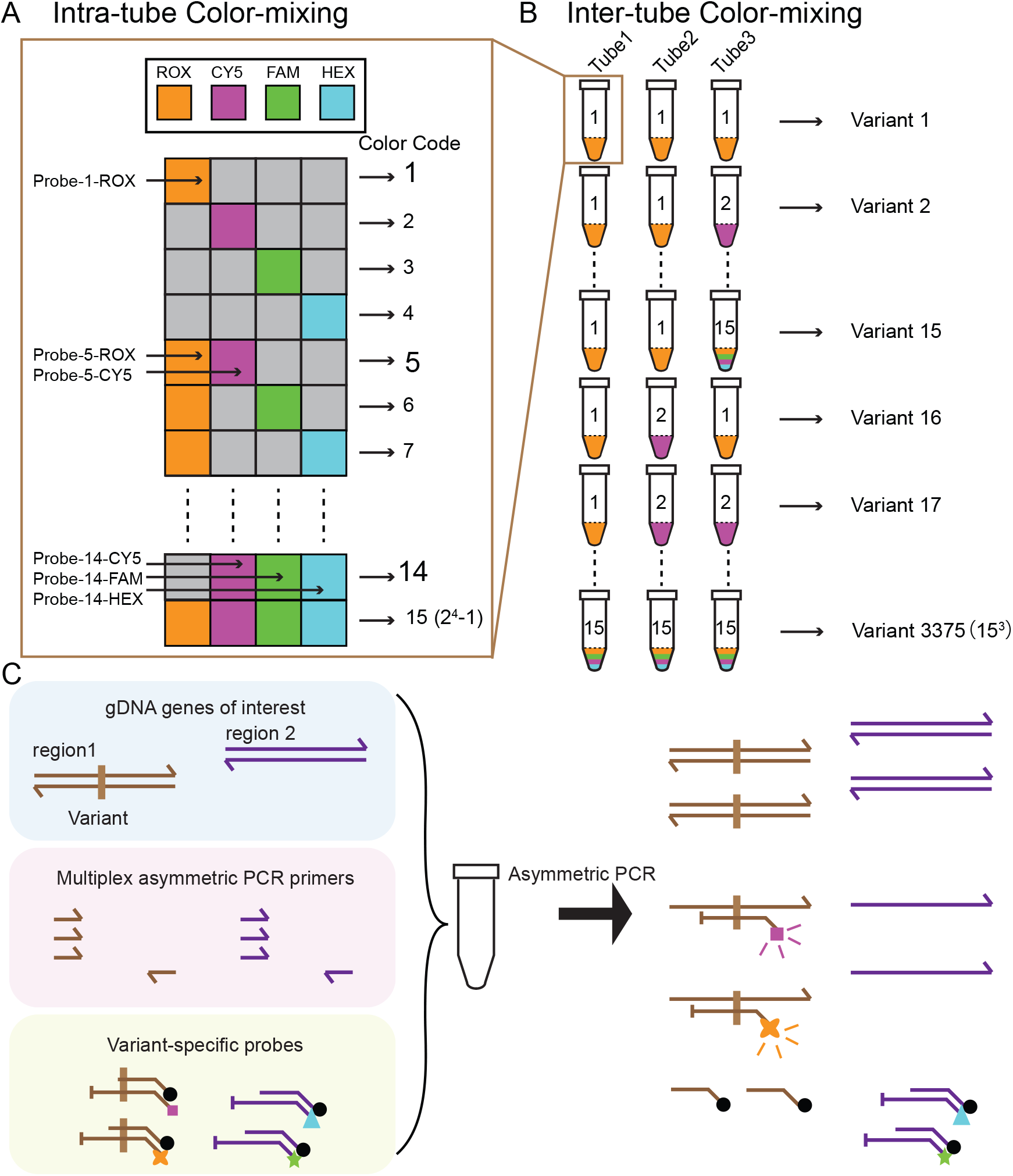
Color-mixing concept and experimental workflow. (A) Intra-tube color mixing. ROX, CY5, FAM, and HEX fluorescence channels are used for each tube. Within one tube, each fluorescence channel’s signal can be identified separately by the instrument, and the final output from each fluorescence channel has 2 states: ‘On’ and ‘Off’. A combination of 4 channels yields 15 (2^<sup>4</sup>^-1) state combinations, and each state combination can be denoted as a color code, with 15 color codes representing 15 individual variants. (B) Inter-tube color-mixing. When increasing the number of tubes, the number of detected variants can be exponentially increased: a combination of 3 tubes can be used to identify as many as 3375 (15^<sup>3</sup>^) variants. (C) Example of experiment workflow. An asymmetric PCR reaction was conducted on mixture of DNA template, multiplex asymmetric primers, and variant-specific toehold probes. Fluorescence signals in different channels were measured before and after asymmetric PCR as background signal and raw signal. The signal differences (raw signal – background signal) were used for evaluating detection performance. After asymmetric PCR, variant-specific toehold probes will bind to the variant amplicons but not to the WT amplicons.

### Inter-tube color-mixing design

The number of simultaneously identifiable targeted variants is exponentially increased when the number of tubes used in an assay increases; the number of targeted variants is 225 (15^2^) for 2 tubes and 3375 (15^3^) for 3 tubes. For example, the color codes ‘1,1,1’ can indicate the presence of variant 1, the color codes ‘1,2,2’ can be used to denote the presence of variant 17, and the color codes ‘15,15,15’ can be used to denote the presence of variant 3375.

### Experimental workflow

The experimental workflow is shown in Figure 1C. The total volume of each reaction was 30 μl (Table. S1 and Table. S2). All multiplex asymmetric primers, a set of rationally designed probes, DNA template, and iTaq Universal Probes Supermix (Bio-Rad) were mixed, followed by execution of a thermal cycling program. The same amount of DNA template was used as the input to each tube. To correct for background noise variation, the fluorescence of the first 8 cycles was measured as the background signal before asymmetric PCR. The asymmetric PCR started with 3 min at 95 °C for polymerase activation, followed by 50 cycles of 30 s at 95 °C for DNA denaturing, 30 s at 64 °C for annealing, and 30 s or 2 min at 72 °C for extension (with the extension time dependent on amplicon length). After the asymmetric PCR, the entire system was heated to 95 °C for 5 min to dissociate all double strands, then cooled to 40 °C for probes binding to the target amplicon. Twenty cycles of fluorescence were measured at 40 °C as the raw signal (Details of protocol are shown in Table. S3 and Table. S4). For further data analysis, the final signals were obtained as raw signal minus background signal for each channel (Fig. S1 and Fig. S2).

### Toehold Probe Design

#### Variant probe design

The design principle for a variant probe is shown in Figure 2A. A variant probe comprises a protector strand (P) and a complementary strand (C)[4, 5].

**Figure 2.**
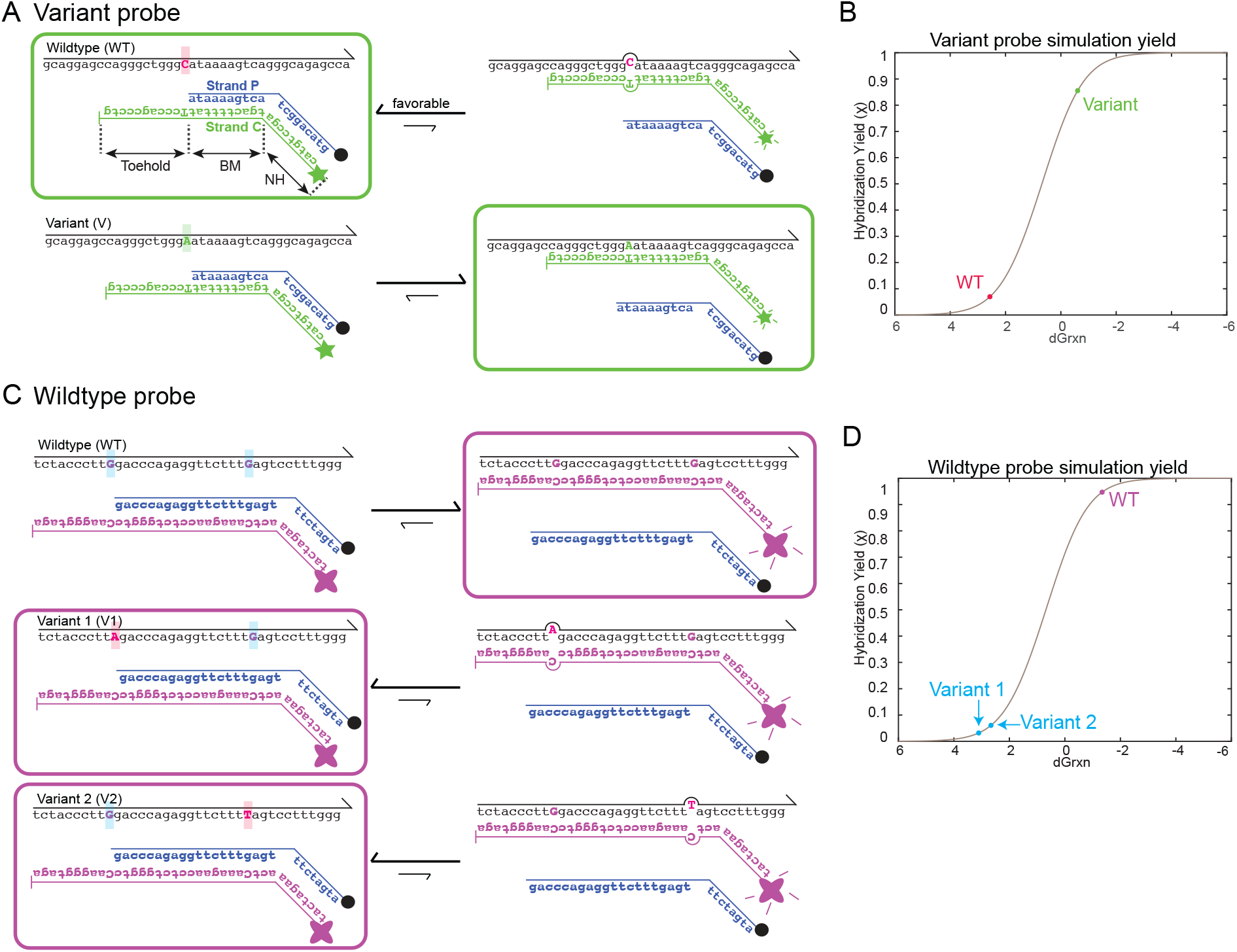
Toehold probe design. (A) The variant probe comprises 2 strands, a complementary strand (C strand) and a protector strand (P strand). The variant probe C strand is designed to be perfectly complementary to a variant strand (V strand), with a fluorophore attached at its 5′ end and a C3 spacer at its 3′ end; the P strand has a quencher at its 3′ end. Green rectangles outline the predominant species at equilibrium: when the V strand is present, the P strand will be displaced, the C strand will bind to the V strand and fluoresce, and the predominant species will be VC; whereas, when the wildtype strand (WT strand) is present, because of the mismatch energy ΔΔG_Mismatch_ between the WT strand and the C strand, the P strand and C strand will favorably bind to each other, the predominant species will be PC, and the fluorophore will be quenched. (B) The simulated hybridization yield of a variant toehold probe binding to a variant strand or a WT strand. Here, 5x V or WT strand, 1xP, 1xPC, and [PC]_0_ = 20 nM were used for simulation. Simulated discrimination factor Q of 32 variant toehold probes is shown in Fig. S6. (C) Wildtype probe design. A wildtype probe also comprises a C strand and a P strand. However, instead of perfectly binding to a specific variant strand, the wildtype probe C strand is designed to perfectly bind to a Wildtype strand (WT strand). The wildtype probe P strand will be displaced by the WT strand and fluoresce. Any variant within the designed region in a wildtype probe will cause a mismatch energy ΔΔG_Mismatch_ to the binding of the C strand, therefore the wildtype probe will not fluoresce when variant amplicons exist. Purple rectangles outline the predominant species of WT toehold probe presented with either WT strand or 2 different variant strands at equilibrium. (D) The simulated hybridization yield of a WT toehold probe binding to variant or WT strand, 5x V or WT, 1xP, 1xPC, and [PC]_0_ = 20 nM. Two different variant strands with different ΔΔG_Mismatch_ will cause slight difference in yield.

When there is no target, the P and C strands will hybridize to form a double-stranded probe.

When a variant amplicon strand (V strand) is present, the standard free energy of variant probe binding to the V strand can be calculated as:

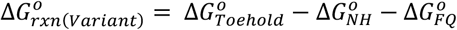

When a WT amplicon strand (WT strand) is present, the standard free energy of variant probe binding to the WT strand can be calculated as:

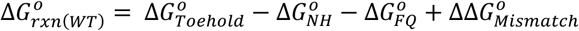

When the variant probe reacts with V strand, the 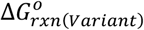 is designed to be below zero; the P strand will be displaced by the V strand, and the C strand will bind to the V strand and fluoresce. When the probe reacts with the WT strand, the 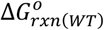 will be higher than 0, and thus will be less thermodynamically favorable; the P and C strands will bind to each other and result in fluorescence quenching. The difference between 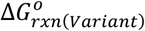 and 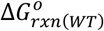 will result in a difference of hybridization yield (Fig. 2B).

Each toehold probe has three regions: the toehold region (Toehold), the branch migration region (BM), and the non-homologous region (NH) (see Fig. 2A). Both Toehold and BM can be rationally designed to perfectly bind to variant amplicons (Supplementary Section 2). The Toehold and BM regions of the C strand are the reverse complement of the V strand, with a fluorophore attached at 5′ end and a C3 spacer at 3′ end. The C3 spacer at 3′ end of the C strand prevents polymerase extension and exonuclease activity during the PCR reaction and therefore functions as a primer and ensures that the probes’ concentrations remain constant after the PCR reaction. The BM region of the P strand is identical to the variant sequence and has a quencher modification at its 3′ end. The NH regions of the P and C strands are random-generated sequences that are designed *not* to be complementary to the amplicon, ensuring that the overall reaction energy will be below 0 kcal mol^-1^ when binding to variant strand and above 0 kcal mol^-1^ when binding to WT strand. Fluorophores and quenchers are attached in the NH region. The empirical energy penalty 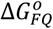 of different fluorophore and quencher binding is different as specified in Supplementary Section 2.

#### Wildtype probe design

Unlike the C strand of a variant probe that perfectly binds to a V strand, the C strand of a wildtype probe is designed to be perfectly reverse complementary to the WT strand (Fig. 2C). When the WT strand is present, the P strand will be displaced by the WT strand, and the C strand will fluoresce. When the V strand is present, with variants at different positions indicated as Variant 1 (V1) and Variant 2 (V2), the P strand binds to the C strand and is thus quenched, resulting in a difference in hybridization yield (Fig. 2D). The fluorescence signal from a wildtype probe will have an ‘On’ state when binding to the WT strand, and an ‘Off ‘ state when binding to the Variant 1or Variant 2 strand.

#### Oligonucleotides and repository samples

All the primers, probes, and synthetic DNA templates were purchased from Integrated DNA Technologies. Solutions of DNA oligonucleotides were stored at 4 °C. Human cell-line gDNA samples (NA18562) were purchased from Coriell Biorepository and stored at −20 °C. The gDNA samples were mixed with synthetic DNA templates at a 1:1 ratio to create samples containing 50% variant sequence. Dilution of gDNA samples and synthetic DNA templates were made in 1× TE buffer with 0.1% Tween 20 from Sigma Aldrich. For the tuberculosis color-mixing validation, with the assumption that clinically extracted genomic DNA of *Mycobacterium tuberculosis* from sputum specimen is low in amount and also contains some human cells or human genomic DNA, the WT sample was a mixture of 2 ng/μL of human cell-line gDNA 18572 (Coriell Biorepository) and 1,000 copies/μL of H37Rv wildtype *Mycobacterium tuberculosis* genomic DNA (ATCC). Each 50% synthetic variant sample was a mixture of 2 ng/μL human cell-line gDNA NA18572 (Coriell Biorepository), 50 copies/μL of wildtype *Mycobacterium tuberculosis* genomic DNA (ATCC), and 50 copies/μL of synthetic template. All templates were prepared in 1× TE buffer with 0.1% Tween 20 (Sigma Aldrich).

## RESULTS

### Identification of pathogenic variants in thalassemia

Thalassemia is a potentially lethal autosomal recessive inherited blood disorder characterized by abnormal synthesis of hemoglobin. There are two types, α- and β-thalassemia; α-thalassemia is involved with genetic pathogenic variants in the HBA1 and HBA2 genes that encode hemoglobin subunit α, while β*-*thalassemia is involved with variants in the HBB gene that encodes hemoglobin subunit β [11]. Individuals with either *α-* or *β-*thalassemia have shortened life expectancy. Therefore, identification of biallelic pathogenic variants in HBA1, HBA2, and HBB may be informative in genotype screening for expectant parents.

We implemented the color-mixing strategy to detect 21 common pathogenic variants in the HBA2 and HBB genes [12]. The detailed design of different variants with their corresponding reference SNP (rs) numbers is shown in Table. S6. We designed a 6-tube assay, with tubes 1 to 3 containing a total of 32 variant probes to identify 21 pathogenic variants and tubes 4 to 6 containing a total of 11 WT probes to further distinguish the samples as homozygous (100%) or heterozygous (50%). Four fluorescence signals were collected from each tube, and a combined 24 fluorescent signals were collected from 6 tubes.

In each tube, each fluorescence channel has an independent threshold to make a call of an ‘On’ or ‘Off’ state. To determine the threshold, we used synthetic DNA with variant strands as positive controls and wildtype genomic DNA as negative controls (Table. S5). Positive controls can be detected by specific variant probes, while negative controls do not include any mutations that variant probes can detect. Using an ROC curve fit, the threshold was determined as the fluorescence value that could best distinguish the final signal (raw – background) of the positive controls from that of negative controls (Fig. S3, Fig. S4). Final signals above threshold are identified as ‘On’ (1), and final signals below threshold are identified as ‘Off’ (0). A sequence of 12 numbers, all 0 or 1, indicate the state combinations used to represent the 21 pathogenic variants.

As an example, the identification of variant 16 is shown in Fig. 3A. In tube 1, the final HEX signal of the variant 16 sample was above the threshold, while the final ROX, CY5, and FAM signals of the variant 16 samples were below the threshold. Therefore, a code of ‘0001’ was generated from tube 1. Likewise, codes of ‘1000’ and ‘0000’ were generated in tubes 2 and 3, respectively. (We only implemented 3 channels in our design for tube 3, therefore only CY5, FAM and HEX signals are shown.) A combination of ‘0001’, ‘1000’ and ‘0000’ from 3 tubes denotes the identification of variant 16.

**Figure 3:**
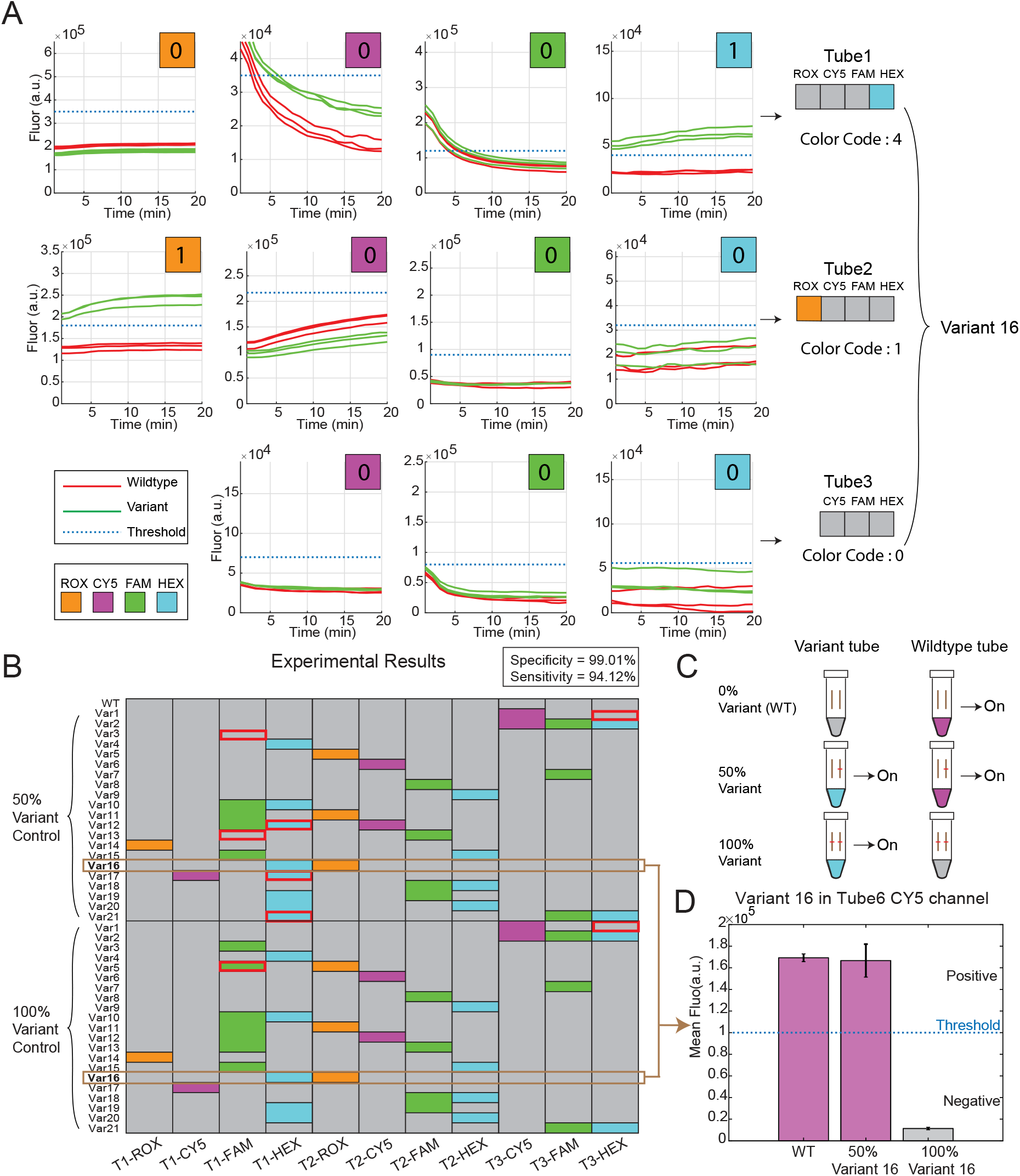
Experimental results of color-mixing detection of pathogenic variants in thalassemia. (A) An example of generating color code for positive control variant 16 in tubes 1 to 3. Triplicate final signals of wildtype sample are displayed as red lines, and the final signals of the variant sample are displayed as green lines. Thresholds are shown as blue dashed lines. In tube 1, only the HEX signal was in ‘On’ state (1), while in ROX, CY5, and FAM, because there were no probes targeting variant 16, the signals of these 3 channels were ‘Off’ (0); a color code of ‘4’ representing ‘0001’ was thus generated in tube 1. Likewise, in tube 2, the ROX signal was ‘On’, while the other channels were ‘Off’. A color code 1 representing ‘1000’ was generated in tube 2. In tube 3, because there was no probe targeting variant 16, the color code of this tube is 0. As we did not include any probe with ROX fluorophore in tube 3, ROX channel’s plot is not shown here. The color codes generated from the 3 tubes identify variant-16. (B) Experimental results of color-mixing design in identifying 21 variants in 3 variant tubes. Different color codes were used to indicate 21 different variants. Probes were used qualitatively; therefore, the color code for synthetic 100% and 50% variant-positive controls are the same. Experimental data resulted in a specificity of 99.01% and a sensitivity of 94.12% when compared with the expected results shown in Supplementary Section 4. (C) Both 50% and 100% variant controls had the same ‘On’ state and color codes in variant tubes. However, 50% variant controls had the ‘On’ state in WT tube, and 100% variant control had the ‘Off’ state in WT tube. (D) The 0%, 50%, and 100% controls of variant 16 can be separated by WT probes in tube 6. Both 0% and 50% control were in ‘On’ state, while the 100% control was in ‘Off’ state.

In Fig. 3B, we show the experimental results of this design for 32 variant probes implemented in tubes 1 to 3 to identify the 21 pathogenic variants in HBA2 gene and HBB gene. Each variant can be denoted by a unique color code; the 50% and 100% synthetic variant controls have the same color code as a result of our qualitative end-point detection approach, and the WT sample is denoted by ‘0000’, ‘0000’ and ‘000’ in tubes 1-3, respectively.

When compared with our expected results (Fig. S5), the experimental results have a sensitivity of 94.12% and a specificity of 99.01%. The few signals that were misread in the FAM and HEX channels in tube 1 indicated the need for further optimization in probe sequence design or probe concentration to achieve the optimal distinction between variant and WT samples.

The 50% and 100% control samples containing the same variant were represented by the same color codes in tubes 1 to 3; we further designed tubes 4 to 6 with wildtype probes to separate the 50% and 100% control samples (Table. S7). For example, 50% and 100% variant 16 were already detected by tubes 1 to 3. And in tube 6, a wildtype probe attached with the CY5 fluorophore was used to separate the 50% and 100% variant-16 control samples (Fig. 3C). WT (0%) and 50% variant samples were in the ‘On’ state, while only the 100% variant sample was in the ‘Off’ state, in accordance with our design (Fig. S7).

Together from the 6 tubes, we are able to separate 21 pathogenic variants related with thalassemia (tubes 1 to 3), and we can also identify individual sample to be heterozygous (50%) or homozygous (100%) (tubes 4 to 6). In tubes 4 to 6, experimental results had a specificity of 99.46% and a sensitivity of 95.24% comparing with the expected results (Fig. S8).

### Thalassemia commercial sample results

We further validated the thalassemia color-mixing design on 4 pre-identified commercial samples. Sample ID, Variant ID in our design, sample name and the rs number are shown in Fig. 4A. Here we only show the results from tubes 1-3 because these pre-identified samples were all heterozygous, therefore the wildtype probes all have a high fluorescence signal in tubes 4-6. The expected fluorescence signals of the 4 samples in tubes 1-3 are also shown here. The 4 samples were experimentally detected and validated and are consistent with our expectation (Fig. 4B, Fig. S9).

**Figure 4:**
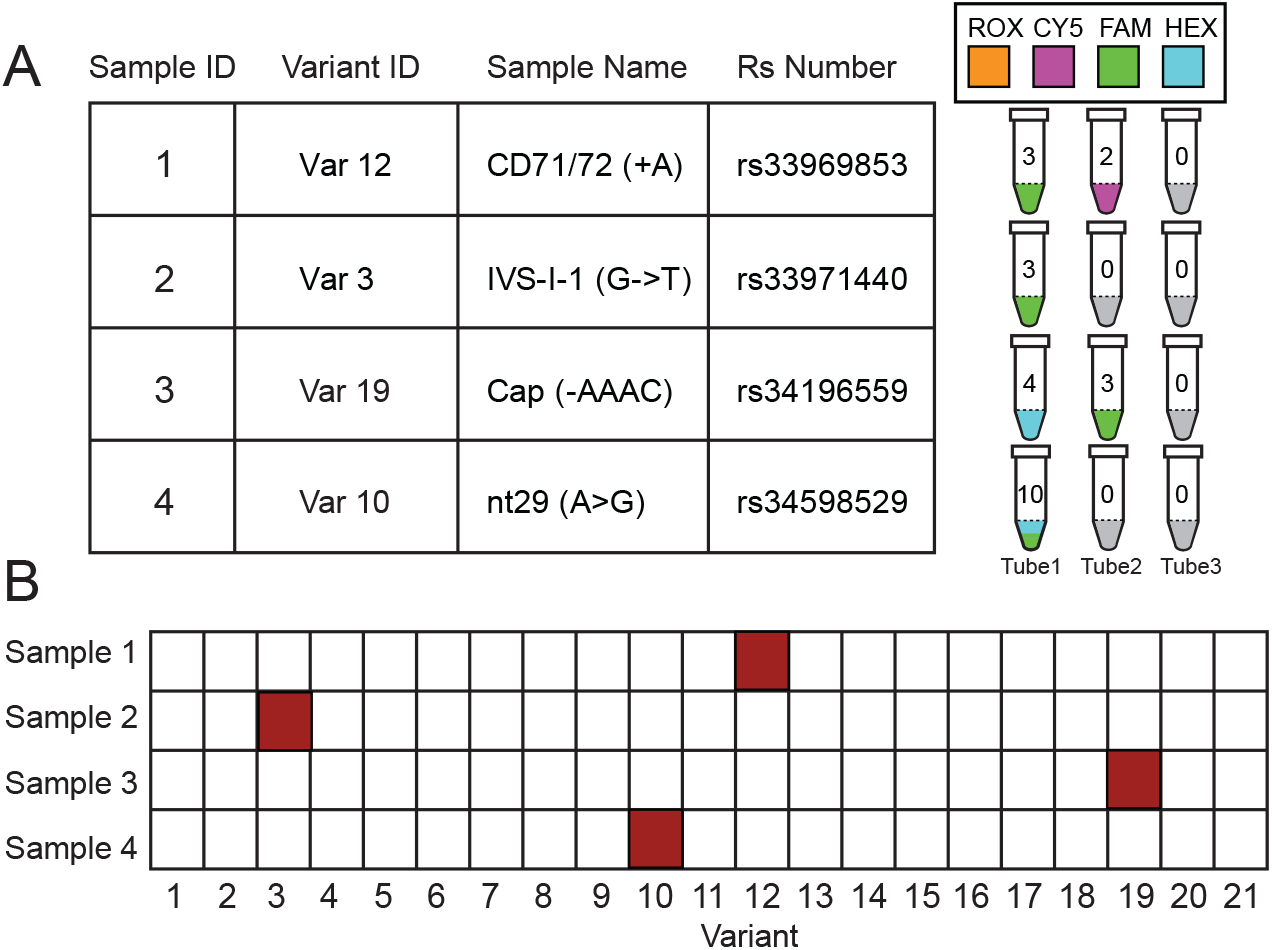
Validation of the thalassemia assay on commercial samples. (A) We tested this assay on 4 commercial samples, and show the sample name, targeted SNP ID, and rs number for our design. On the right, the expected fluorescence for our color-mixing design is shown. Noted that the variants for these four samples were pre-identified, and all the samples were heterozygous samples. (B) The call of variants in 4 samples are shown; all samples were successfully identified.

### Drug resistance in tuberculosis

We further implemented the color-mixing strategy in the detection of drug resistance in tuberculosis (TB). Tuberculosis, usually caused by *Mycobacterium tuberculosis* bacteria (MTB), is one of the 10 deadliest contagious diseases worldwide. According to the World Health Organization (WHO) Global Tuberculosis Report 2019, an estimated 10 million people fell ill with TB in 2018. Successful pharmacologic treatment began in the 1940s, when streptomycin use was found effective. Soon after that, it became clear that single-drug therapy led to the evolution of drug-resistant MTB strains, and therefore high failure rates in TB treatment [16].

Multi-drug-resistant TB (MDR-TB) is resistant to the two most potent anti-TB drugs, rifampicin (RIF) and isoniazid (INH). Approximately 5% of newly diagnosed TB cases are rifampicin-resistant TB (RR-TB), and 78% of the 5% RR-TB are multi-drug-resistant TB (MDR-TB). Thus, RR-TB and MDR-TB are of significant concern at the global level, indicating the urgency for development of detection methods for drug-resistance mutations [17]. Prompt diagnosis of MDR-TB would tell clinicians to switch to a second-line drug regimen such as fuoroquinolones [17].

Several mutations in specific genes across the bacterial genome are known to lead to the drug resistance of MTB (Fig. 5A), so it would help to identify drug resistance-related mutations among the hot spot genetic regions in the MTB genome.

**Figure 5:**
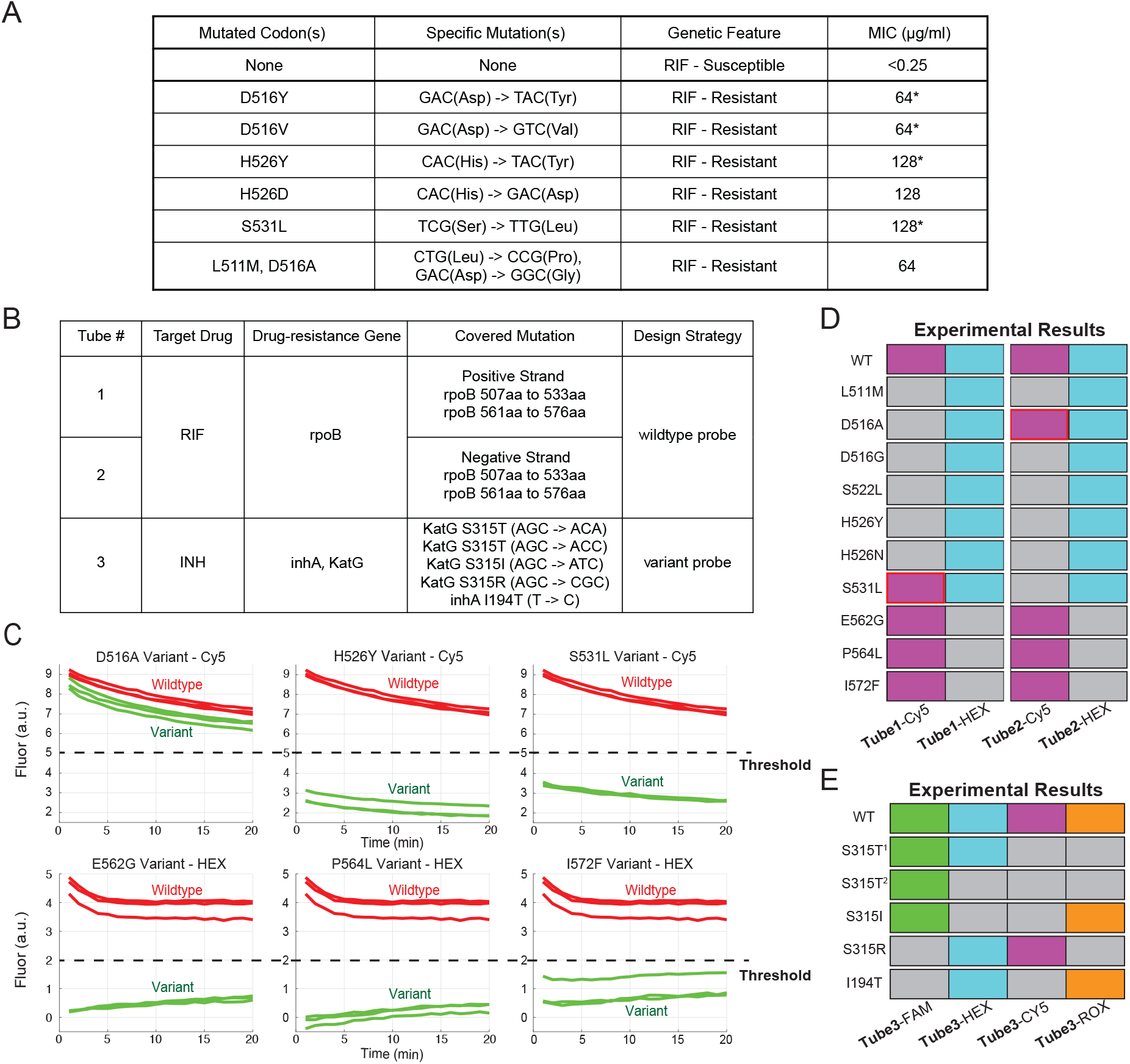
Color-mixing strategy in detecting TB-drug-resistance mutations in *Mycobacterium tuberculosis*. (A) Drug resistance in *Mycobacterium tuberculosis*. Drug resistance, i.e., the lack of drug susceptibility of MTB to the anti-tuberculosis drug, is shown in the difference of minimal inhibitory concentrations (MIC) between each clinical strain, which is a quantitative parameter describing the concentration of antibiotic to which a bacterium is sensitive. Here, taking the RIF MIC as an example, the RIF MIC of the susceptible strain (non-RIF-resistant strain) is different from the RIF MICs of mutant strains [13]. (B) Summary of assays. Using wildtype probes, tubes 1 and 2 target the RRDR and Cluster II regions of the *rpoB* gene, whose mutations cause the RIF-resistance of MTB. Tube 3 targets multiple mutations in the INH resistance-related *katG* and *inhA* genes using variant probes. (C) In each reaction, 20 ng of human DNA + 1000 MTB DNA copies (either 100% genomic DNA or 50% variant template and 50% genomic DNA) were used as input. A final readout was obtained by comparing the final fluorescence with threshold. All wildtype sample final signals are displayed as red lines, all variant sample final signals are displayed as green lines, and thresholds based on the ROC curve for each channel are displayed as dashed lines. For example, variant-D516A is called wildtype in tube 2 because its final fluorescence is above threshold and variant-P564L is called variant in tube 2 because its final fluorescence is below threshold; summaries of all the tubes are shown in 5d and 5e. (D) Comparison between expected results and experimental results in wildtype tubes. In tubes 1 and 2, the H37Rv wildtype genomic DNA and 10 synthetic variant templates were tested and summarized. (E) Comparison between expected and experimental results in tube 3. In tube 3, the H37Rv wildtype genomic DNA and 5 synthetic variant templates were tested and summarized. **Notes:** MIC: minimal inhibitory concentration, the lowest concentration of a chemical, usually a drug, that prevents visible growth of a bacterium or bacteria. *when there were multiple strains with the same mutation under MIC test, the results shown are the median MICs. S315T^1^: S315T(AGC → ACA) S315T^2^: S315T(AGC → ACC)

Strategies have been applied to detect MTB drug-resistance mutations before. Culture-based phenotypic drug susceptibility test (DST) methods are currently the gold standard for drug resistance detection, but these methods are time-consuming (usually from 2 weeks to 6 weeks) and require sophisticated laboratory infrastructure, qualified staff, and strict quality control [28]. Many other methods have been reported before, e.g., multiplex PCR [18, 19], melting temperature (Tm) analysis in the real-time PCR reaction [20], microarray [21], and real-time PCR using molecular beacon probes [22, 23]. However, these technologies are usually very complex to operate, not user-friendly, too time-consuming, or too expensive due to the reagents or instruments. Here we applied the color-mixing strategy to detect drug resistance mutations in TB, achieving rapid and simultaneous detection of drug resistance mutations in multiple genes and regions within 3.5 hours.

We designed a 3-tube assay covering the RIF-resistance gene *rpoB* and the INH-resistance genes *katG* and *inhA* (Fig. 5B). *RpoB* encodes the β subunit of RNA polymerase. An 81-bp region of the *rpoB* gene has been reported to be the rifampin resistance-determining region (RRDR) [24]. Although around 90% - 96% [24, 25] of RIF resistance in MTB is associated with mutations inside the RRDR region, there is another region in *rpoB* called the cluster II region [25, 26] that contains the remaining 4% - 10% [24, 25] of RIF-resistance mutations in MTB.

There are difficulties in detecting drug-resistance mutations in the *rpoB* gene. 1) Previous methods predominantly covered the RRDR region, resulting in lower sensitivity and specificity. 2) The 81-bp RRDR and 48-bp Cluster II-related regions are relatively long and high in GC content, but also rich in drug-resistance mutations. Most previous methods were limited to detecting only a few hot spot mutations, leading to incomplete mutation profiling and less accurate diagnostic information. To solve these difficulties, we designed an assay to cover both the RRDR and cluster II regions within the *rpoB* gene in both positive and negative strands (tubes 1 and 2 in Fig. 5B) using wildtype probes to achieve comprehensive detection of *rpoB* drug-resistance mutations.

INH resistance commonly occurs when mutations occur in the *katG* gene or the regulatory regions of *inhA* gene. *KatG* functions to encode catalase-peroxidase, which is an enzyme that can activate INH. Mutations that occur in the *katG* gene, especially at codon 315, have a high correlation with INH resistance. Another target of active INH is nicotinamide adenine dinucleotide-dependent enoyl-acyl carrier protein reductase, which is encoded by the *inhA* gene [27]. We designed a panel (shown as tube 3 in Fig. 5B) to simultaneously detect multiple hot spot mutations within one tube in this work. The targets of tube 3 are mainly single-nucleotide variants and relatively shorter than those in tubes 1 and 2; thus, targeting single mutations in only positive or negative strands using variant probes is sufficient for sensitivity and specificity.

Each of tubes 1 and 2 contains 2 WT probes to identify any drug-resistance mutation occurring within the target region. Only signals from the Cy5 and HEX fluorescence channels were collected from each tube. Examples of the identification of some variants in tube 2 are shown in Fig. 5C. Each experiment was performed in triplicate. In Fig. 5C, the final signals of variants other than the D516A variant are below the wildtype samples as well as the threshold. That means that all variants except the D516A variant could be confidently detected in tube 2. The threshold is determined by the ROC curve as described before. The experimental results of our design for tube 1 and tube 2 are shown in Fig. 5D. In tubes 1 and 2, if the final signal of one unknown sample is above the threshold, it indicates that the sample does not have mutations in the targeted region, otherwise the sample should contain drug-resistance mutations in the targeted region. Ideally, every variant regardless of mutation pattern or strand could be detected in each tube. S531L could not be distinguished with wildtype in tube 1, and D516A is indistinguishable from wildtype in tube 2, marked as red frames in Fig. 5D. More details are discussed in the Supplementary Section 7, the failure of D516A is probably from the minimum difference of standard free energy between the binding of the probe to the variant and to the WT as shown in Table. S8 and Fig. S10. The strong hairpin structure formed around S531L probably lead to the in-distinguishable between S531 and the WT in tube 1 (Fig. S11). The failures in detecting S531L in tube 1 and D516A in tube 2 imply the necessity of targeting both the positive and negative strands. In these cases, combining the results of tubes 1 and 2 would help provide a more accurate and comprehensive result because S531L is easily separated with WT in tube 2 and the final signal of D516A is much lower than that of WT in tube 1.

Tube 3 contains 10 variant probes to identify 5 drug-resistance mutations, and 4 fluorescence signals would be collected. The experimental and expected results for tube 3 are shown in Fig. 5E and Fig. S12. Each variant is denoted by a unique color code, as shown in Fig. S12. We tested all the covered mutations in tube 3: one variant (I194T) in *inhA* and four variants (S315T^1^, S315T^2^, S315I and S315R) in *katG* (Table. S9). Each variant could be distinguished from WT and other variants with 100% sensitivity as shown in Fig. 5E.

In this manner, we successfully demonstrated the use of the color-mixing strategy to simultaneously detect drug-resistance mutations of MTB.

## DISCUSSION

In this study, we present a color-mixing strategy that utilizes end-point fluorescence detection to achieve highly multiplex probing of genomic variants simultaneously. The color-mixing strategy involves two steps: first, enrichment of regions of interest by an asymmetric PCR, and second, detection of genetic variants by a set of rationally designed multiplex toehold probes. A set of toehold probes were either all variant probes or all wildtype probes. We implemented the color-mixing strategy into two applications, the detection of thalassemia-related pathogenic variants and the detection of drug-resistance mutations in tuberculosis. The results demonstrate that a simple color-mixing strategy can be adapted to different applications for different purposes, and that it can be used to identify variant and/or wildtype sequences.

Thalassemia is a potentially fatal autosomal recessive blood disorder. We designed a 6-tube assay that included 3 variant tubes to identify 21 pathogenic variants in HBA2 and HBB genes, and another 3 wildtype tubes that can distinguish the synthetic 50% and 100% variant samples that we designed to mimic the clinical heterozygous and homozygous samples. We note that heterozygous samples are much more clinically common than homozygous samples because the homozygous variant may be lethal in the fetus. Our design can be used for prenatal screening for pathogenic variants and can therefore provide information for expectant parents.

We also designed a 3-tube assay to detect drug-resistance mutations in MTB. Our designed assay includes two WT tubes that can identify any mutation in both an 81-bp region and a 48-bp region that are related to RIF resistance. It includes another variant tube that can identify five mutations previously reported to be highly correlated with INH resistance. Using the designed multiplex wildtype probes, we experimentally validated the identification of 10 synthetic variants within two target regions in the *rpoB* gene. Each template used in the experiments was designed to mimic a clinical sputum sample extracted from the lung, with a mixture of bacteria and human cells. We validated tube 3 (variant tube) by probing for all the variants with 100% sensitivity. Some loci or some mutations may be difficult to be detected either because the sequences are high in GC content and may form secondary structures, or because the energy penalty brought by the mutation is barely distinguishable by probes (Supplementary Section 7). Our design can guide clinicians to decide specific treatment plans for each patient promptly.

The color-mixing strategy shows a way to use a qPCR instrument to detect multiple variants. We believe that this method is a general method for detection of multiple variants in other diseases beyond thalassemia and tuberculosis. This method can be used for detection in clinical samples in the future. With limited access to thalassemia and tuberculosis clinical samples, we could not verify the method in actual clinical samples, but hope to find collaborators for such verification in the near future.

## ACKNOWLEDGEMENTS

The authors thank Paul Dolber for editorial assistance.

## AUTHOR CONTRIBUTIONS

NGX, KZ, PS, and DYZ conceived the project. NGX, KZ, and PS performed the experiments and analyzed the data. RL and JL performed experiments for Thalassemia commercial samples. NGX, KZ, PS and DYZ wrote the manuscript with input from all authors.

## ADDITIONAL INFORMATION

Correspondence may be addressed to DYZ. DYZ declares a competing interest in the form of consulting for and significant equity ownership in NuProbe Global, Torus Biosystems and Pana Bio.

## DATA AVAILABILITY

The sequences of the DNA oligos and the oligo concentrations used for fluorescence experiments are included as an Excel file accompanying this manuscript. The fluorescence data from our experiments are in Supplementary Information.

## CODE AVAILABILITY

The MATLAB code used for the DNA sequence design and fluorescence data analysis is available in the Supplementary Information.

