## Supplementary file for "High-throughput variant detection using a color-mixing strategy"

**Section 1. Experimental Protocol**

**Oligonucleotides and repository samples.** All the primers, probes, and synthetic DNA templates were purchased from Integrated DNA Technologies (IDT). Primers were purchased as standard desalted DNA oligonucleotides and synthetic templates as desalted double strand fragments (gBlocks). Solutions of DNA oligonucleotides were stored at 4 °C. In thalassemia color-mixing validation, Human cell-line gDNA sample NA18562 (Coriell Biorepository) was stored at −20 °C. The gDNA samples were mixed with synthetic DNA templates at a 1:1 ratio to create samples containing 50% variant sequence. Dilution of gDNA samples and synthetic DNA templates were made in 1× TE buffer with 0.1% Tween 20 from Sigma Aldrich.

In tuberculosis color-mixing validation, with the assumption that clinically extracted genomic DNA of *Mycobacterium tuberculosis* from sputum specimen is low in amount and also contains some human cells or human genomic DNA contamination, the WT sample was the mixture of 2ng/ul human cell-line gDNA 18572 (Coriell Biorepository) and 1,000 copies/ul H37Rv wildtype *Mycobacterium tuberculosis* genomic DNA (ATCC), each 50% synthetic variant sample was prepared to 2ng/ul human cell-line gDNA NA18572 (Coriell Biorepository), 50 copies/ul wildtype *Mycobacterium tuberculosis* genomic DNA (ATCC) and 50 copies/ul synthetic template. All templates were prepared in 1×TE buffer with 0.1% Tween 20 from Sigma Aldrich.

**Thermal Cycling protocol.** As shown in Table S1 and Table S2. The reaction mixture of asymmetric PCR primers mixture, multiplex toehold probes mixture, template solution and iTaq polymerase Supermix were pooled in a 0.2mL Eppendorf PCR tube. Detailed concentration of Primer Mix and Probe Mix in each tube is provided in Supplementary Excel.

| Reagent | Add in Volume (μL) |
|---|---|
| iTaq universal probes supermix | 50 |
| Tube specific Primer Mix | 14 |
| Tube specific Probe Mix | 10 |
| Milli-Q water | 16 |
| Template solution | 10 |
| Total Volume | 100 |

Table S1. Reaction mixture formulation for Thalassemia color-mixing design.

| Reagent | Add in Volume (μL) |
|---|---|
| iTaq universal probes supermix | 50 |
| Tube specific Primer Mix | 10 |
| Tube specific Probe Mix | 10 |
| Milli-Q water | 20 |
| Template solution | 10 |
| Total Volume | 100 |

Table S2. Reaction mixture formulation for tuberculosis color-mixing design.

The reaction mixture was triplicated to 96-well PCR plate with each well 30 μL of the reaction mixture. The 96-well PCR plate was put into an Applied Biosystem Quantstudio Flex 7 or Biorad CFX96 instrument. Thermal cycling program is shown in Table S3 and S4.

| Step | Temperature | Duration | Signal Collection |
|---|---|---|---|
| 1. Background Signal Measurement | 40°C | 1 min | Yes |
| Repeat Step 1 for 8 times | | | |
| 3. Denaturation | 95°C | 3 min | |
| 4. Denaturation | 95°C | 30 s | |
| 5. Annealing | 64°C | 30 s | |
| 6. Extension | 72°C | 2 min | |
| Repeat Step 4 - 6 for 50 times | | | |
| 10. Denaturation | 95°C | 5 min | |
| 11. Raw Signal measurement | 40°C | 1 min | Yes |
| Repeat Step 11 for 20 times | | | |

Table S3. Thermal cycling reaction protocol for thalassemia color-mixing design and tube 1 and tube 2 of tuberculosis color-mixing design.

| Step | Temperature | Duration | Signal Collection |
|---|---|---|---|
| 1. Background Signal Measurement | 40°C | 1 min | Yes |
| Repeat Step 1 for 8 times | | | |
| 3. Denaturation | 95°C | 3 min | |
| 4. Denaturation | 95°C | 30 s | |
| 5. Annealing | 64°C | 30 s | |
| 6. Extension | 72°C | 30 s | |
| Repeat Step 4 - 6 for 50 times | | | |
| 10. Denaturation | 95°C | 5 min | |
| 11. Raw Signal measurement | 40°C | 1 min | Yes |
| Repeat Step 11 for 20 times | | | |

Table S4. Thermal cycling reaction protocol for tube 3 of tuberculosis color-mixing design.

**Section 2. Introduction to Toehold probe design**:
**2.1 Variant Probe**

The design of Variant probes in thalassemia comprising 8 steps. All steps were implemented by MATLAB.

1. Identify the genomic variant of interest. We downloaded the downloaded the reference genomic sequence of HBA2 and HBB gene and *Mycobacterium tuberculosis* H37Rv from NCBI. We did literature research to locate Variant of interest on the reference sequence. Variant of interest was located on the reference sequence by their rs number. For single variant, WT sequences and variant sequences were marked and recorded for future design. Noted that in our design, the variants included SNP and INDEL.
2. Decide to use positive (+) strand or negative (-) strand as target amplicon. We decided to use positive strand or negative strand based on the mismatch ddG between the WT sequence and its corresponding variant sequence on the positive strand and negative strand. Our goal was trying to minimize the binding of WT template to the Variant probe, which means the selection of the larger mismatch ddG, so that the difference of standard free energy between the probe react to the variant and the probe react to the WT is higher.
   For example, 'aCc' > 'aAc' in positive strand is corresponding to 'gGt' > 'gTt' in negative strand. According to our calculation based on published parameters[1], at 40°C and a salinity of 0.18:
   $$ddG_{Mismatch}\,(\text{'aCc'} > \text{'aAc'}) = 3.53,$$
   $$ddG_{Mismatch}\,(\text{'gGt'} > \text{'gTt'}) = 1.54.$$
   Therefore, we decided to target the 'aCc' > 'aAc' in positive strand.
3. Determine Design requirements. We designed the free standard energy of Toehold region, BM region and NH region based on our empirical values. To specify, at 40°C and a salinity of 0.18, the $dG_{Toehold}$ was designed to be around -9.5 kcal/mol, the $dG_{BM}$ to be around -6 kcal/mol and $dG_{NH}$ to be around -5.5 kcal/mol, note that $dG_{NH}$ didn't include the binding energy of fluorophore and quencher $dG_{F\text{-}Q}$, which is usually around -3.5~-4 kcal/mol. To specify for tuberculosis design, the $dG_{Toehold}$ was designed to be around -9 kcal/mol, the $dG_{BM}$ to be around -16 kcal/mol, though to optimize the performance, we modified individual probes to be around -12 kcal/mol.
4. Design the BM region. BM region was first designed with the base of interest in the middle or on the side, and the $dG_{BM}$ to be around -6 kcal/mol as mentioned in Step 3.
5. Design the Toehold region. We then designed Toehold region to be next to the BM region, with a $dG_{Toehold}$ to be around -9.5 kcal/mol as mentioned in Step 3.
6. Design the NH region. We then designed the NH region on the other side of the BM region by first generating a random sequence and select a random portion of the sequence that looks sufficiently dissimilar to the target sequence. And the $dG_{NH}$ to be around -5.5 kcal/mol. The empirical energy penalty $dG_{(FQ)}$ of fluorophore and quencher binding is different among different fluorophores and to be more specific, for the FAM, the $dG_{(FQ)}$

[1] SantaLucia, J., Jr, & Hicks, D. (2004). The thermodynamics of DNA structural motifs. *Annual review of biophysics and biomolecular structure*, *33*, 415–440. https://doi.org/10.1146/annurev.biophys.32.110601.141800

is from -3.5 to -4 kcal/mol; for the HEX, it is -4 kcal/mol; for the CY5, it is -4 kcal/mol; for the ROX, it is from -3.4 to -4 kcal/mol.

7. Piece together Quencher Sequence. Quencher sequence was finalized by piece the sequence of BM and NH regions together.
8. Piece together Fluorophore Sequence. Fluorophore sequence was finalized by piece the reverse complementary sequence of NH, BM and Toehold regions together.

### 2.2.1 Wildtype probe - thalassemia

For thalassemia color-mixing design, WT probe is almost the same as the Variant probe, except that in Step 4, the BM region should be the same in length and sequence as WT target amplicon, instead of Variant target amplicon. Noted that unlike the Variant probe to be specific to one variant, WT probe can perfectly bind to the WT target amplicon, and any variant amplicon within the region covered by the probe will not favorably react with the WT probe. For tuberculosis design, the BM region covered all the sequence as WT target amplicon, thus the BM region of Wildtype probe for tuberculosis is long due to the lengthy target region of *Mycobacterium tuberculosis*.

**Section 3. Data analysis pipeline**

**Background Noise correction.** As there's variation of background fluorescence signal in each well of the qPCR instrument. To correct this background noise, we collect the fluorescence signal twice: the first time before the asymmetric PCR reaction, and the signal was termed as Background Signal (BS) (Fig.S1), the second time after the asymmetric PCR reaction, and was termed as Raw Signal (RS). We did background correction to generated Final Signal (FS) by subtracting BS from FS (Fig.S2).

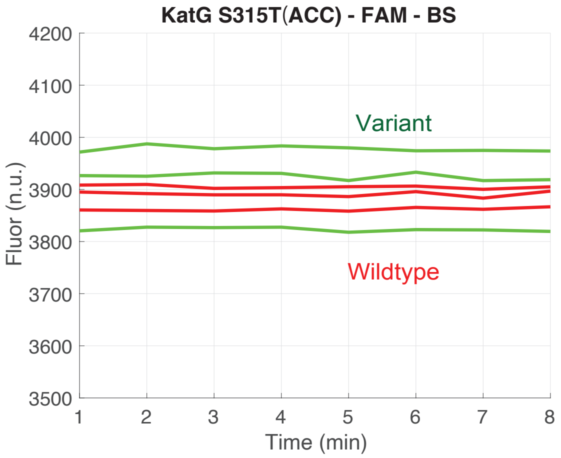


Fig. S1. Example of Background Signal (BS)

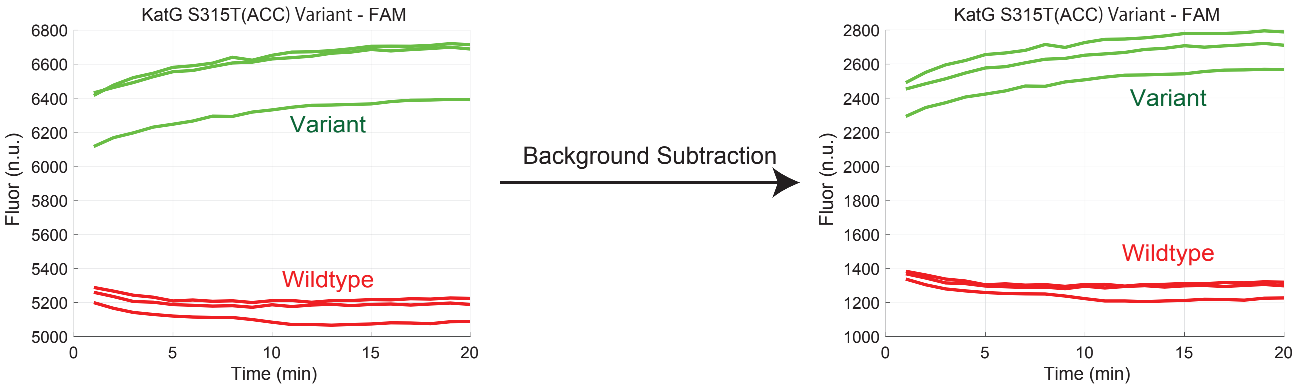


Fig. S2. Example of background noise subtraction

**Control Sample preparations.** 43 control samples including 0%, 50 % and 100% variant samples were prepared. 21 homozygous samples were prepared by serial dilution from the synthetic gBlocks (IDT) using 1xTE 0.1% Tween. 50 % variant samples were prepared by mixing the diluted homozygous samples and the NA18562 at 1:1 ratio. Control 1 is the Wildtype NA18562, control 2-22 were 50% variant samples and control 23-43 were 100% variant samples. Control ID, variant ID and variant ratio are shown in Table S5.

| Control ID | variant ID | Variant Ratio | Control ID | variant ID | Variant Ratio |
|---|---|---|---|---|---|
| 1 | Wildtype | 0% | | | |
| 2 | Variant 1 | 50% | 23 | Variant 1 | 100% |
| 3 | Variant 2 | 50% | 24 | Variant 2 | 100% |
| 4 | Variant 3 | 50% | 25 | Variant 3 | 100% |
| 5 | Variant 4 | 50% | 26 | Variant 4 | 100% |
| 6 | Variant 5 | 50% | 27 | Variant 5 | 100% |
| 7 | Variant 6 | 50% | 28 | Variant 6 | 100% |
| 8 | Variant 7 | 50% | 29 | Variant 7 | 100% |
| 9 | Variant 8 | 50% | 30 | Variant 8 | 100% |
| 10 | Variant 9 | 50% | 31 | Variant 9 | 100% |
| 11 | Variant 10 | 50% | 32 | Variant 10 | 100% |
| 12 | Variant 11 | 50% | 33 | Variant 11 | 100% |
| 13 | Variant 12 | 50% | 34 | Variant 12 | 100% |
| 14 | Variant 13 | 50% | 35 | Variant 13 | 100% |
| 15 | Variant 14 | 50% | 36 | Variant 14 | 100% |
| 16 | Variant 15 | 50% | 37 | Variant 15 | 100% |
| 17 | Variant 16 | 50% | 38 | Variant 16 | 100% |
| 18 | Variant 17 | 50% | 39 | Variant 17 | 100% |
| 19 | Variant 18 | 50% | 40 | Variant 18 | 100% |
| 20 | Variant 19 | 50% | 41 | Variant 19 | 100% |
| 21 | Variant 20 | 50% | 42 | Variant 20 | 100% |
| 22 | Variant 21 | 50% | 43 | Variant 21 | 100% |

Table S5. Control Sample Information

**Threshold Selection.** As shown in Table S5, we prepared 0%, 50% and 100% variant controls. Each control can be categorized to be positive and negative for each channel in one tube. For example, in tube 1, only a ROX probe was designed to target the Variant 14, and because control 15 contains 50% Variant 14 and control 36 contains 100% Variant 14. Therefore, control 15 and 36 were categorized as positive in ROX channel in tube1, while the rest 41 controls were categorized as negative. Positive controls were colored in purple and negative controls were colored in blue in Fig S3a. X axis is the 43 controls, and y axis is the background fluorescence corrected Final Signal.

We show here an example ROC fit to find the optimized Threshold in ROX channel in tube 1 (Fig S3b) in thalassemia color-mixing assay. The Threshold was selected from between the minimum and maximum of the Final signal and was determined to maximize the AUC. When the Threshold was set at 3*10^5, all the positive controls can be successfully distinguished from negative controls. Final signals of all 4 channels in 3 tubes were plotted in Fig S4.

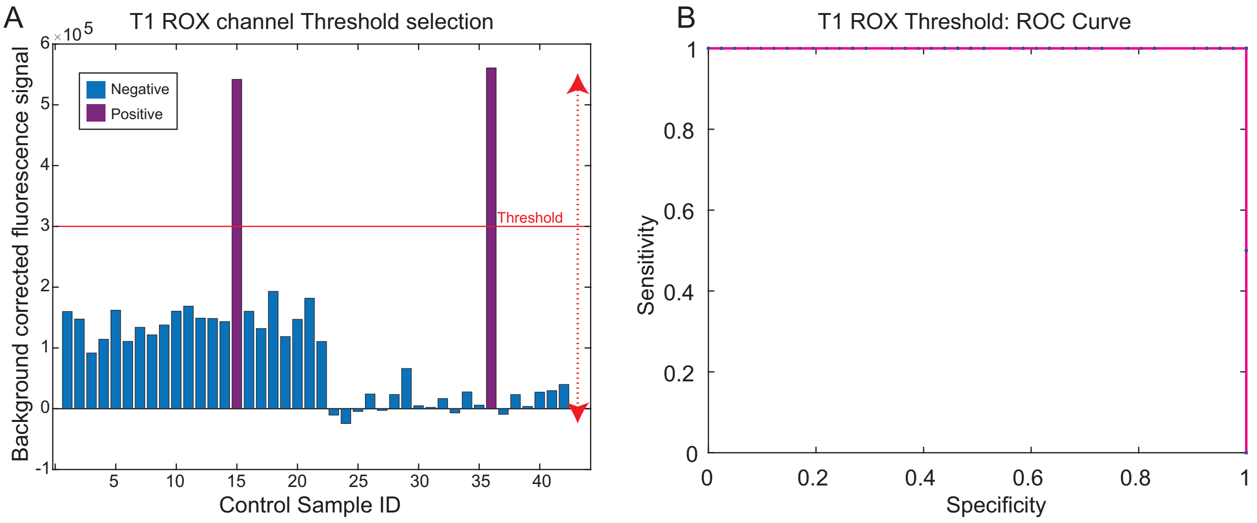


Figure S3. Threshold selection in tube 1 ROX channel

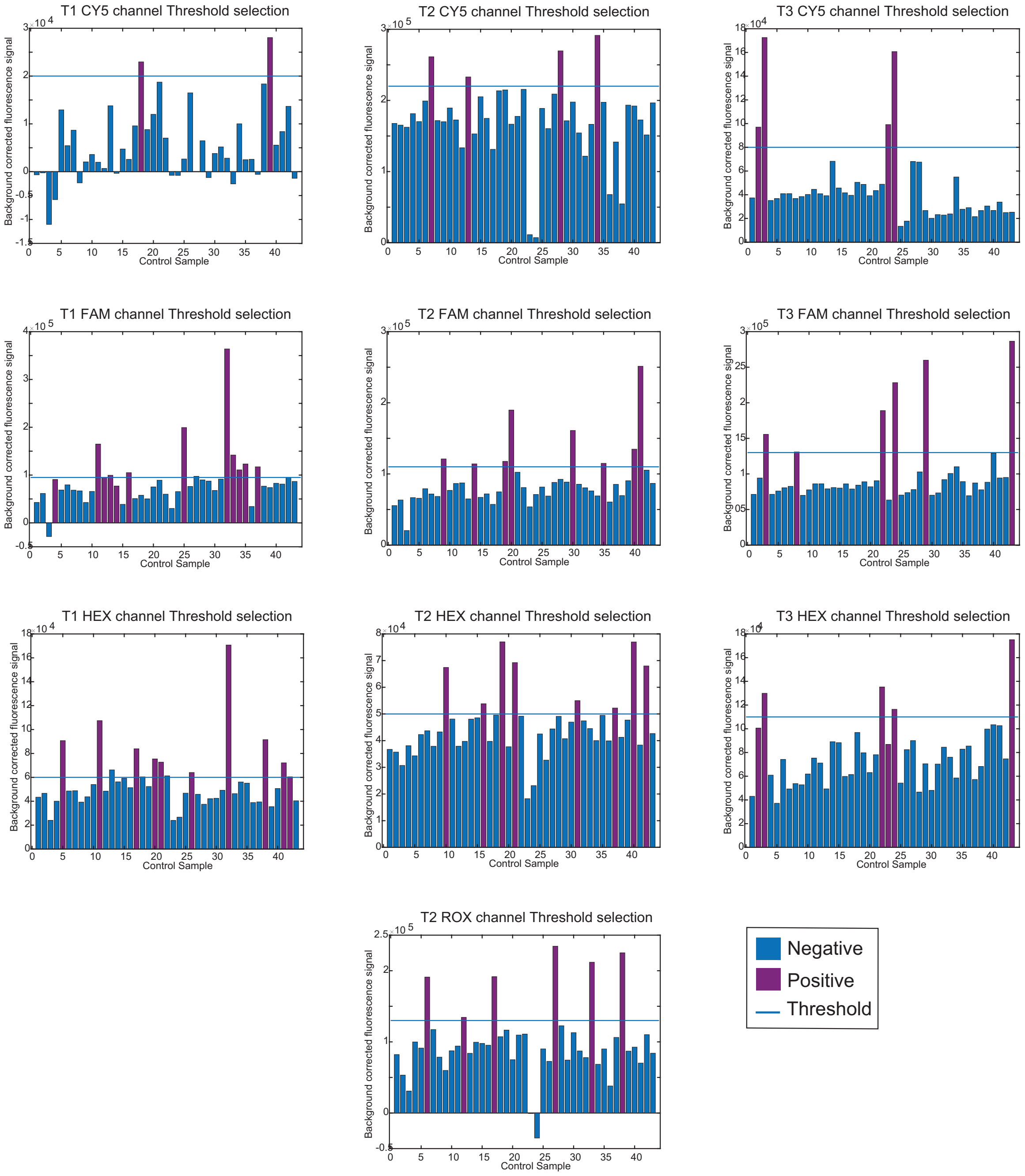


Figure S4. Threshold selection in tube 1- tube 3 in 4 fluorescence channels

**Section 4. Detailed design of color mixing in thalassemia tube 1 to 3**

In our implementation of color mixing strategy to detect 21 pathogenic variants in thalassemia, we specified Variant ID, gene name, dbSNP ID, HGVS name, position and mutation to use in Table S6. Noted that the mutation to use were based on the preference of more significant $\Delta\Delta G_{Mismatch}$ as mentioned in Supplementary Section 2. Frequent pathogenic SNPs and INDELs are included in 21 variants, of which 2 are on HBA2 gene and 19 are on the HBB gene.

| Variant ID | Gene | dbSNP ID | HGVS name | Position | mutation to use |
|---|---|---|---|---|---|
| 1 | HBA2 | rs41397847 | c.377T>C | 173548 | T>C |
| 2 | HBA2 | rs41464951 | c.427T>C | 173598 | T>C |
| 3 | HBB | rs33971440 | c.92+1G>T | 5226929 | C>A |
| 4 | HBB | rs35684407 | c.91A>G | 5226931 | T>C |
| 5 | HBB | rs35383398 | c.45_46insG | 5226977 | dup |
| 6 | HBB | rs33931746 | c.-78A>G | 5227099 | T>C |
| 7 | HBB | rs33991059 | c.113G>A | 5226779 | C>T |
| 8 | HBB | rs33915217 | c.92+5G>C | 5226925 | C>G |
| 9 | HBB | rs35532010 | c.84_85insC | 5226939 | dupG |
| 10 | HBB | rs34598529 | c.-79A>G | 5227100 | T>C |
| 11 | HBB | rs35099082 | c.315+5G>C | 5226572 | C>G |
| 12 | HBB | rs33969853 | c.216_217insA | 5226675 | dup |
| 13 | HBB | rs63749977 | c.94delC | 5226798 | delG |
| 14 | HBB | rs34451549 | c.316-197C>T | 5225923 | G>A |
| 15 | HBB | rs33922842 | c.130G>T | 5226762 | C>A |
| 16 | HBB | rs34500389 | c.-82C>A | 5227103 | G>T |
| 17 | HBB | rs80356821 | c.124_127delTTCT | 5226765_5226768 | delAGAA |
| 18 | HBB | rs33986703 | c.52A>T | 5226970 | T>A |
| 19 | HBB | rs34196559 | c.-11_-8delAAAC | 5227030_5227033 | delTTTG |
| 20 | HBB | rs35755331 | c.126delC | 5226766 | delG |
| 21 | HBB | rs33950507 | c.79G>A | 5226943 | C>T |

Table S6. Genetic variants targeted in the implementation of color mixing for thalassemia

**Simulation of binding yield.** A simulation was conducted to predict the discrimination factor Q of the binding yield difference of 32 variant probes bind to Variant strand and WT strand.

1. The reaction of variant probe binds to the Target strand (T) can be written as follow:

$$T + PC \leftrightarrow TC + P$$

The T strand can be either a Variant strand (V) or a Wildtype strand (WT), and the binding energy of toehold probe to variant strand $\Delta G_{rxn(Variant)}$and to WT strand $\Delta G_{rxn(WT)}$ can be calculated as:

$$\Delta G^{o}_{rxn(Variant)} = \Delta G^{o}_{Toehold} - \Delta G^{o}_{NH} - \Delta G^{o}_{FQ} \quad (1)$$

$$\Delta G^{o}_{rxn(WT)} = \Delta G^{o}_{Toehold} - \Delta G^{o}_{NH} - \Delta G^{o}_{FQ} + \Delta\Delta G^{o}_{Mismath} \quad (2)$$

2. The change in reaction energy $\Delta G^{o}_{rxn}$ is related to equilibrium constant Keq by the equation:

$$\Delta G^{o}_{rxn} = -RT\ln(Keq) \quad (3)$$

Therefore, equilibrium constant $Keq_{(Variant)}$ and $Keq_{(WT)}$ can be calculated from the equation (1)(2) and (3).

3. In our design, we can start with the $[PC]_0 = c$, $[P]_0 = c$, $[T]_0 = ac$. We defined the $x = [TC]\infty$ to be the concentration of hybridization product TC at equilibrium. The equation of Keq can be written as follows:

$$Keq = \frac{[TC]_\infty [P]_\infty}{[T]_\infty [PC]_\infty}$$

$$Keq = \frac{x\,(x + c)}{(ac - x)(c - x)}$$

$$(Keq - 1)x^2 - (Keq * (a - 1)c - c)x + Keq * ac^2 = 0 \quad (4)$$

4. The final concentration of $[TC]_\infty$ at equilibrium therefore can be calculated from equation (4), and if we assume that a > 1, then the binding yield $\chi = \frac{[TC]\infty}{[TC]\infty + [PC]\infty} = \frac{x}{c}$. In the main paper Fig.2B, the $\Delta G^o_{rxn(Variant)}$ = 0.61 kcal/mol at 40 ℃ and a salinity of 0.18, binding yield $\chi =$ 85.6%. And the $\Delta G^o_{rxn(WT)}$ = 2.57 kcal/mol at 40 ℃ and a salinity of 0.18, the binding yield $\chi = 7.0\%$.

5. Discrimination factor was defined as $Q = \frac{\chi_{(Variant)}}{\chi_{(WT)}}$, the Q in Fig.2B is 12.3. The summary of simulated discrimination factor Q of 32 toehold probes in tube 1 to 3 is shown in Fig.S6. In general, small INDEL have a higher Q than the SNV because of stronger $\Delta\Delta G^o_{Mismath}$.

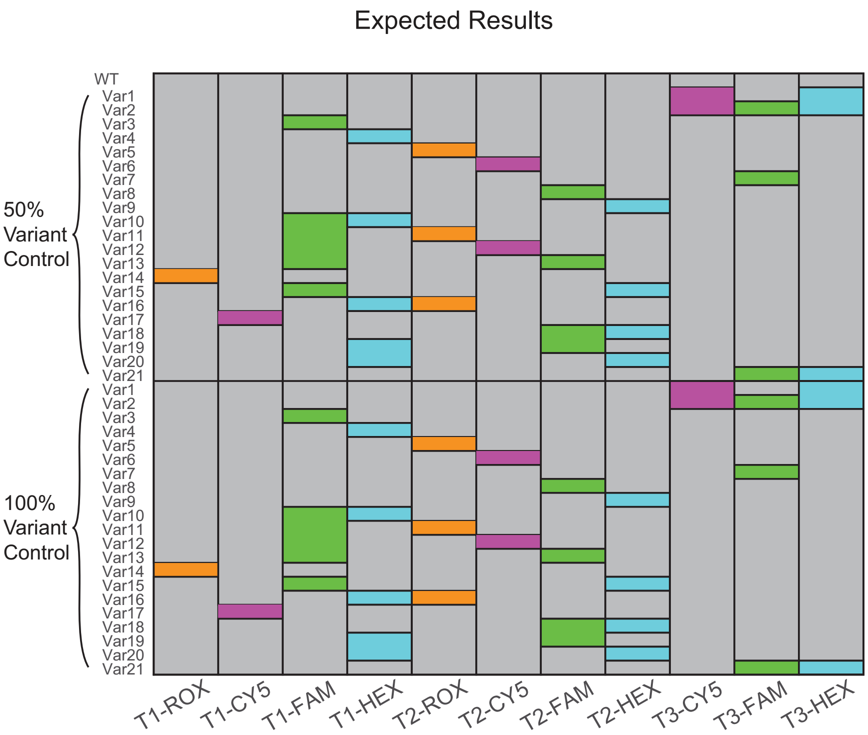


Figure S5. Expected results of thalassemia pathogenic genetic variants detection in tube 1 to tube 3.

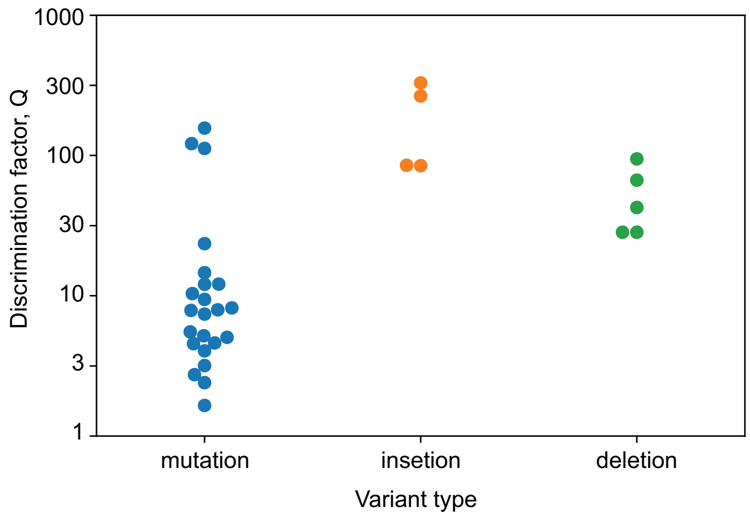

Figure S6. Simulation of discrimination factor Q of 32 toehold probes in tube 1 to 3.

**Section 5. Design and results of color mixing in thalassemia tube 4 to 6**

After the identification of Variants in tube 1 to tube 3. We also implemented 3 Wildtype tubes to separate 100% Variant controls (homozygous variant) and 50% Variant controls (heterozygous variant). Design of color-mixing strategy in WT tubes is shown in Table S5. Unlike the Variant probe, one probe targets one specific variant sequence, a WT probe can cover several variants within a region. For instance, as the variants 7, 15, 17 and 20 are close to each other in the genome, one WT probe can be used to confirm the WT genotype at the region from variant 7 to 20. In this way, if the sample is a 50% variant sample with either of these 4 variants, the WT probe will bind to the wildtype sequence and fluoresce; while if the sample is 100% variant, the WT probe will not favorably bind to the variant sequence, thus won't fluoresce. We therefore separated 21 variants into different groups due to their positions, variants that are close to each other can be covered by the same WT probe. In our design, 11 WT probes attached with different fluorophores in tube 4 to 6 can distinguish between 100% Variant controls and 50% Variant controls.

| Tube 4<br>(+) strand HBA, (+) strand HBB | | Tube 5<br>(-) strand HBB | | Tube 6<br>(-) strand HBB | |
|---|---|---|---|---|---|
| Variants/WT detected | Fluorescence | Variants/WT detected | Fluorescence | Variants/WT detected | Fluorescence |
| WT-1 | ROX | WT-13 | ROX | WT-12 | ROX |
| WT-7-15-17-20 | CY5 | WT-3-4-8-9-21 | CY5 | WT-16-6-10 | CY5 |
| WT-2 | FAM | WT-11 | FAM | WT-14 | FAM |
| WT-5-18 | HEX | | | WT-19 | HEX |

Table S7. WT probes design in tube 4 to 6

Example results of distinguishment of 100% and 50% controls containing the same variant 4 is shown in Fig S7. Subtracted CY5 fluorescence signals of 0%, 50% and 100% Variant controls were plotted in red, blue and green. Threshold determined by ROC fit was plotted in blue dashed line.

According to our design, the WT probe will bind to the WT template and fluoresce, because 0 % and 50% Variant controls both contained the WT template, therefore after the asymmetric PCR, 0% and 50% Variant controls had above-threshold fluorescence signals, while the 100% Variant control didn’t. 50% Variant control had a lower signal than 0% variant control, this is reasonable because the 50% variant control contained less WT molecules than the 0% variant control. Nevertheless, a proper Threshold was determined to just separate the samples contain WT sequence than those not because our assay was designed to be qualitative but not quantitative. Note that we also observed in other channels that the 50% Variant and 0% Variant controls gave approximately the same level of fluorescence.

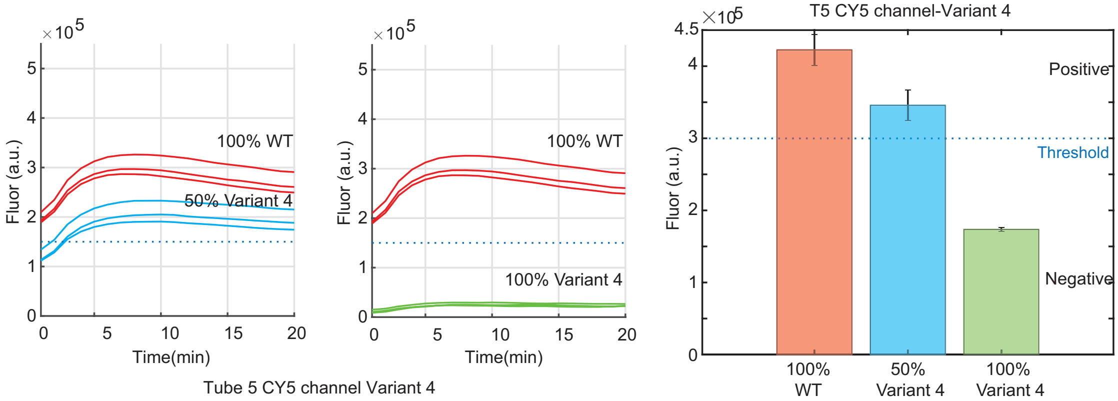


Figure S7. Experimental results of 0%, 50% and 100% Variant 4 in Tube 5 Cy5 Channel

The experimental results of all the 0%, 50% and 100% variant controls are shown in Fig S8. Note that we specified some of the data points to be N/A, it's not that we didn't do the experiments, but the data points couldn't be used as references because of the length limitation of the synthetic gBlocks. All the gBlocks we ordered had a length of 700 bp, therefore gBlocks covering region of interest in HBA2 can't cover regions in HBB genes, and vice versa. For example, Variant 1 is on HBA2 and synthetic gBlocks that mimic Variant 1 can only cover the 700 bp region in HBA2. However, in reality, a sample should contain both the regions in HBA2 and HBB genes. And a sample contains Variant 1 on HBA2 gene can be wildtype in HBB gene. Therefore, a 100% Variant 1 sample can be used as a WT for other positions in HBB gene. Nevertheless, our synthetic Variant 1 sample didn't contain the WT sequence in HBB gene and couldn't be targeted by the WT probe and fluoresce. Therefore, it couldn't be used as a reference in specific channels that probes are targeting HBB gene. We specified these data points in 100% variant controls to be N/A to minimize the error in our Threshold selection and avoid false negative results.

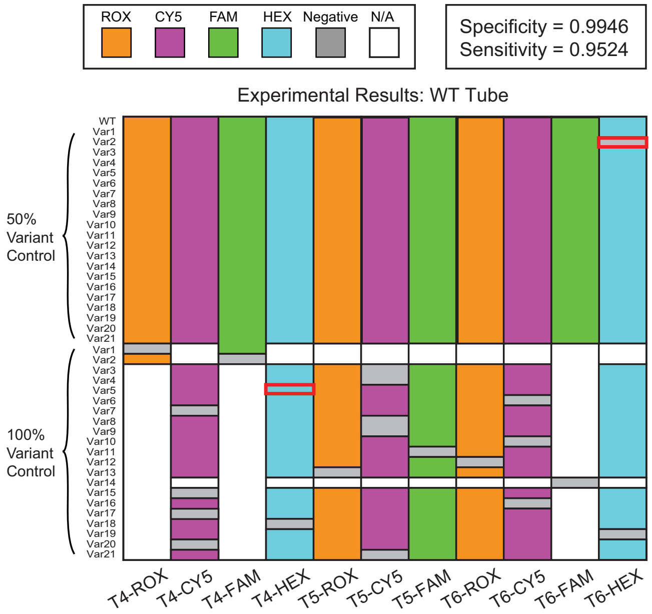


Figure S8. Experimental results of thalassemia pathogenic genetic variants detection in tube 4 to tube 6.

**Section 6. Results of commercial samples**

We validated our assay on four commercial thalassemia samples. Results of four samples are shown in Fig.S9. As the commercial samples are heterozygous samples. We showed here the results of tube 1 to tube 3. X axis is the tube number, y axis is the Signal/Threshold ratio. We analyzed in this way so that if the ratio is above 1, it's in 'On' state, and if the ratio is below zero, it's in 'Off' state. All 4 samples can be accurate identified by our assay. Noted that we did observe some high variation in FAM and HEX channels which was consistent with our positive and negative controls' result. This may indicate that 1. Probe sequences and concentration may need to be optimized to reach the best performance. 2. In general CY5 channel and ROX channel work better than FAM and HEX channel, in the future, we can re-organize the color-mixing design to include more CY5 and ROX probes.

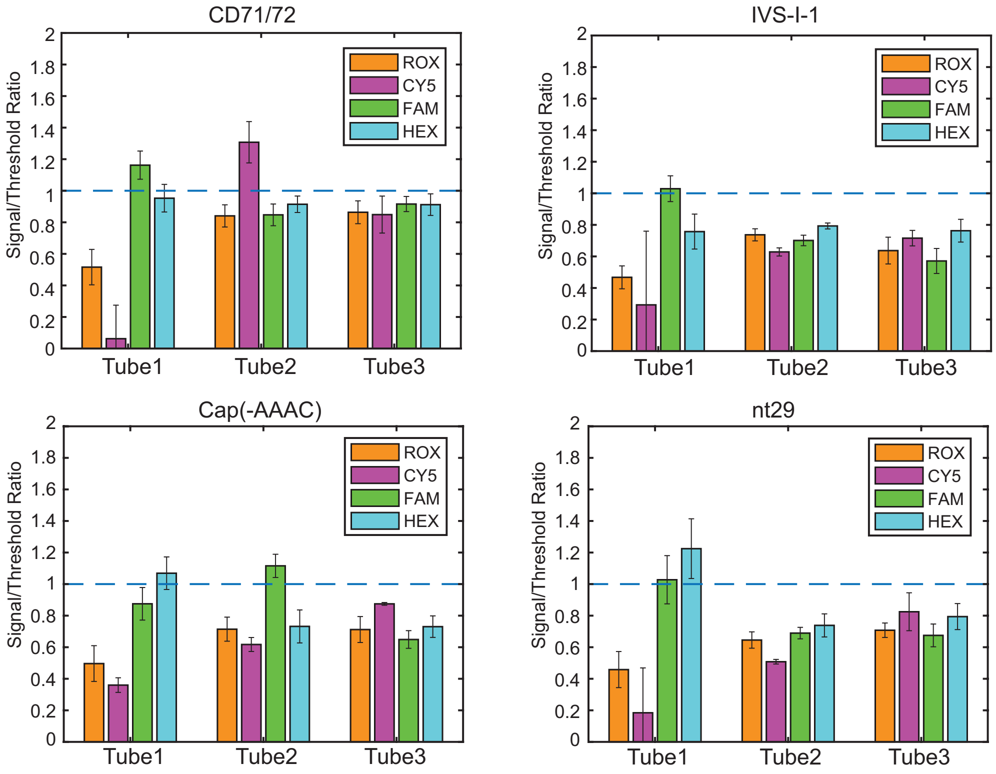


Fig S9. Results of commercial samples

**Section 7. Design details of color mixing in tuberculosis tube 1 to 2**

Experimental results are shown in Fig. 5D, each of tube 1 and tube 2 has one sample that do not meet our experimental expectation (marked as red frame). We explored the mechanism in this section. First of all, we summarized all the ΔΔG of each mutation as shown in Table S8 and Figure S10. We found that the mutation D516 A failed to be detected in tube 2 had the minimum ΔΔG, that means the difference of standard free energy between the probe to D516A and the probe to the WT is much small. That makes mutation D516A is barely distinguishable in tube 2.

| Target mutation | Tube 1 ΔΔG | Tube 2 ΔΔG |
|---|---|---|
| L511M | 2.7212 | 4.3383 |
| D516G | 1.7333 | 3.4815 |
| D516A | 3.8198 | 1.4494 |
| S522M | 2.8587 | 4.9509 |
| H526N | 3.4675 | 4.1546 |
| H526Y | 2.9021 | 4.4473 |
| S531L | 2.8587 | 4.9509 |
| E562G | 2.1857 | 3.8104 |
| P564L | 3.0203 | 4.2443 |
| I572F | 2.7771 | 2.5011 |

Table S8. ΔΔG of each variant used in validation of tube 1 & 2 assay

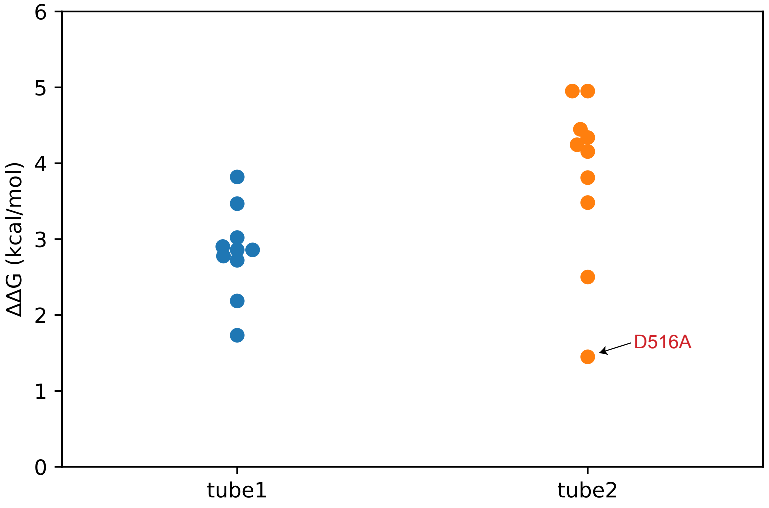


Figure S10. ΔΔG of each variant used in validation of tube 1 & 2 assay

Because the *Mycobacterium tuberculosis* genome is GC rich, so the possibility of forming a secondary structure is high. We used Nupack to check the secondary structures of the amplicons in both tube 1 and tube 2. The red shaded area is what the WT probe with Cy5 modification targets, the blue shaded area is what the WT probe with HEX modification targets. We found that the neighboring region of mutation S531L formed a hairpin with a 7nt stem and 86% GC content. Thus, we hypothesize that the failure of detecting S531L in tube 1 is due to the formation of a high-

GC hairpin within the target region (Fig.S11). The $dG_{Toehold}$ of Cy5-WT probe in tube 2 is stronger than the $dG_{Toehold}$ of Cy5-WT probe in tube 1, stronger $dG_{Toehold}$ makes the WT probe more likely to open the secondary structure. In this way, we think this is why tube 2 is successfully detects S531L but tube 1 not.

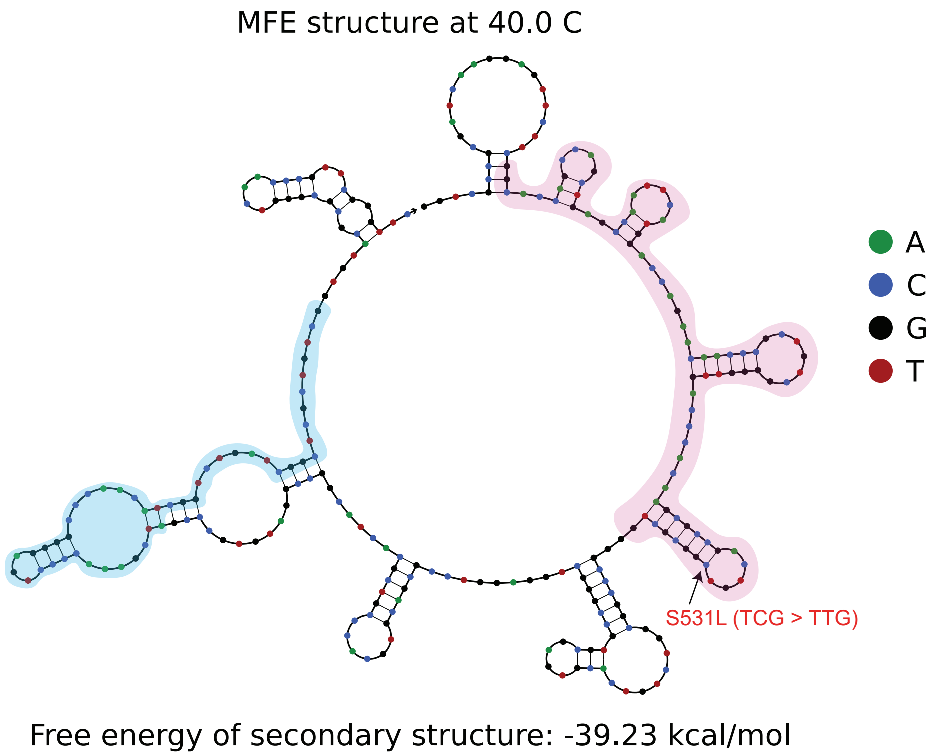


Figure S11. Secondary structure folding of amplicon in tube 1 and tube 2.

**Section 8. Design details of color mixing in tube 3 of tuberculosis**

In our implementation of color mixing strategy to detect 5 INH-resistance variants in *Mycobacterium tuberculosis* (MTB), we specified Variant ID, gene name, position, AA change, mutation to use in Table S9. Noted that the mutation to use were dependent on the preference of more significant $ddG_{Mismatch}$ as mentioned in Supplementary Section 2. Our experimental results (Fig. 5E) match the expected results, shown in Fig. S12.

| Variant ID | Gene | Position | AA change | mutation to use |
|---|---|---|---|---|
| 1 | KatG | 2155167 - 2155168 | S315T | AGC -> ACA |
| 2 | KatG | 2155168 | S315T | AGC -> ACC |
| 3 | KatG | 2155168 | S315I | AGC -> ATC |
| 4 | KatG | 2155169 | S315R | AGC -> CGC |
| 5 | inhA | 1674782 | I194T | ATC -> ACC |

Table S9. Genetic variants targeted in the implementation of color mixing for tuberculosis

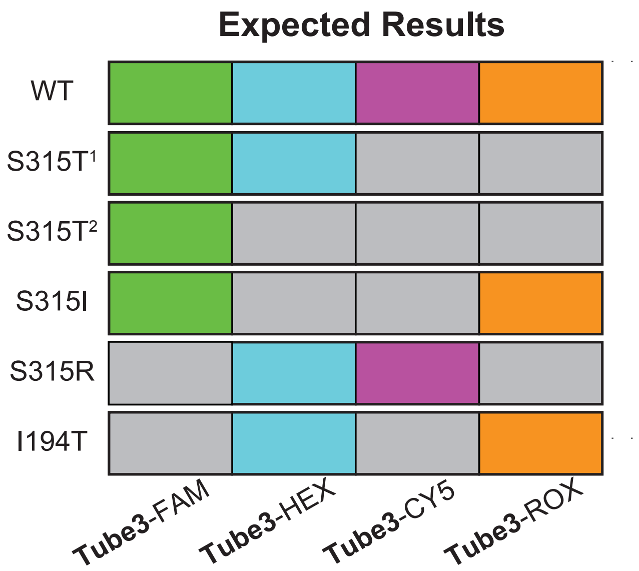


Figure S12. Expected results of tuberculosis drug-resistance variants detection in tube 3.
